# Fiber orientation downsampling compromises the computation of white matter tract-related deformation

**DOI:** 10.1101/2021.12.07.471622

**Authors:** Zhou Zhou, Teng Wang, Daniel Jörgens, Xiaogai Li

## Abstract

Incorporating neuroimaging-revealed structural details into finite element (FE) head models opens vast new opportunities to better understand brain injury mechanisms. Recently, growing efforts have been made to integrate fiber orientation from diffusion tensor imaging (DTI) into FE models to predict white matter (WM) tract-related deformation that is biomechanically characterized by tract-related strains. Commonly used approaches often downsample the spatially enriched fiber orientation to match the FE resolution with one orientation per element (i.e., element-wise orientation implementation). However, the validity of such downsampling operation and corresponding influences on the computed tract-related strains remain elusive. To address this, the current study proposed a new approach to integrate voxel-wise fiber orientation from one DTI atlas (isotropic resolution of 1 mm^3^) into FE models by embedding orientations from multiple voxels within one element (i.e., voxel-wise orientation implementation). By setting the responses revealed by the newly proposed voxel-wise orientation implementation as the reference, we evaluated the reliability of two previous downsampling approaches by examining the downsampled fiber orientation and the computationally predicted tract-related strains secondary to one concussive impact. Two FE models with varying element sizes (i.e., 6.37 ± 1.60 mm and 1.28 ± 0.55 mm, respectively) were incorporated. The results showed that, for the model with a large voxel-mesh resolution mismatch, the downsampled element-wise fiber orientation, with respect to its voxel-wise counterpart, exhibited an absolute deviation over 30° across the WM/gray matter interface and the pons regions. Accordingly, this orientation deviation compromised the computation of tract-related strains with normalized root-mean-square errors up to 30% and underestimated the peak tract-related strains up to 10%. For the other FE model with finer meshes, the downsampling-induced effects were lower, both on the fiber orientation and tract-related strains. Taken together, the voxel-wise orientation implementation is recommended in future studies as it leverages the DTI-delineated fiber orientation to a larger extent than the element-wise orientation implementation. Thus, this study yields novel insights on integrating neuroimaging-revealed fiber orientation into FE models and may better inform the computation of WM tract-related deformation, which are crucial for advancing the etiological understanding and computational predictability of brain injury.

**Graphic abstract:** 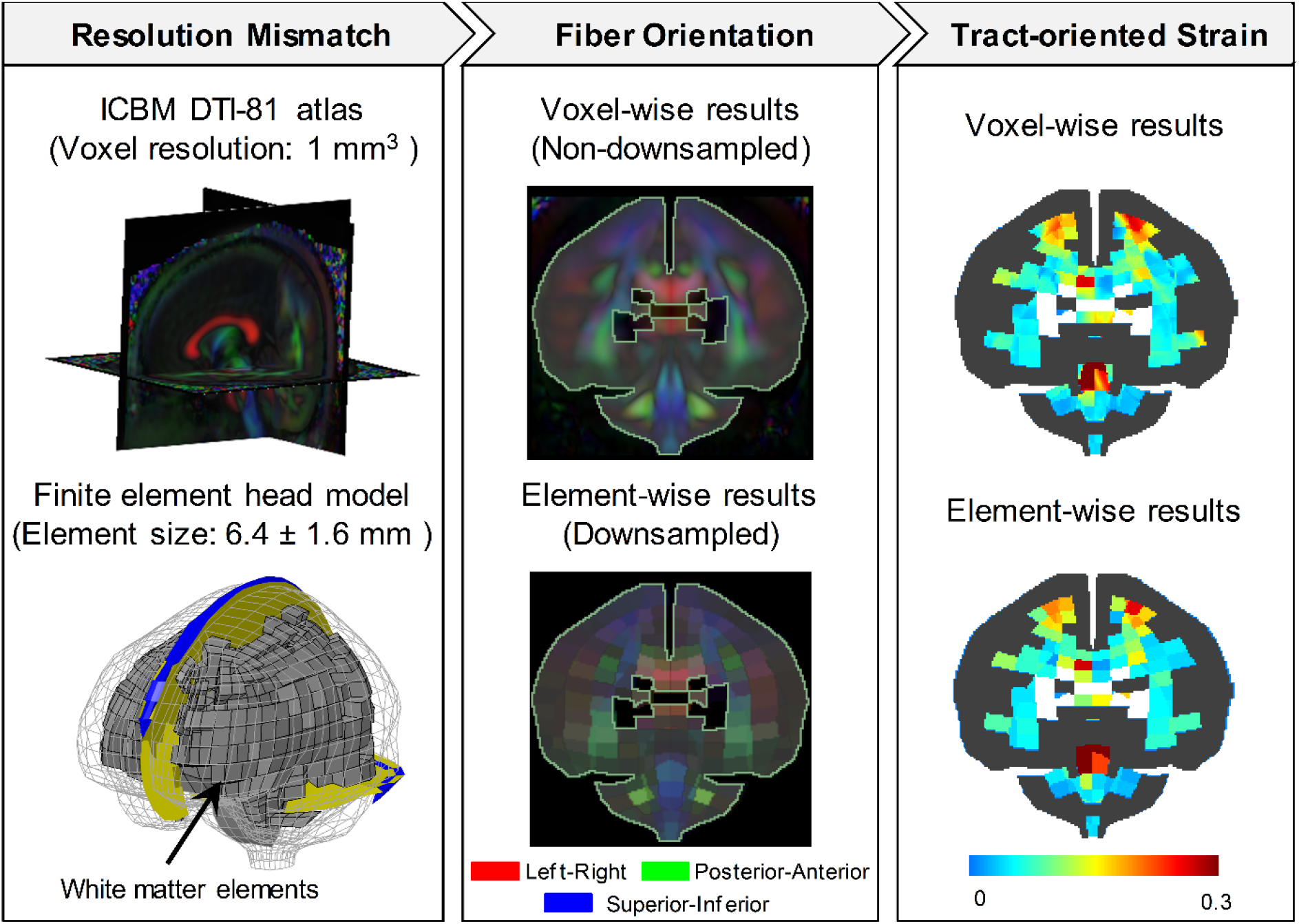

## 1. Introduction

Traumatic brain injury (TBI) is a disruption in the brain’s normal function or other evidence of brain pathology secondary to an external insult, typically in the form of direct head impact and inertial loading (Menon et al., 2010). Depending on the severity of insults, the outcome of TBI may vary from temporary unconsciousness accompanied by transient neuronal circuit dysfunction, e.g., concussion (Wolf et al., 2017), to persistent vegetative state along with profound neuronal apoptosis throughout the white matter (WM), e.g., diffuse axonal injury (Maxwell et al., 2003). Given its high morbidity and mortality and the large number of people affected (Brazinova et al., 2021; Peterson et al., 2019), TBI is a critical public health and socio-economic threat worldwide. Sadly, current prediction, prevention and diagnosis strategies for brain injury are still largely in their infancy due to a lack of fundamental understanding of TBI pathogenesis.

Biomechanical models based on the finite element (FE) method are promising instruments to decipher the physics underlying TBI. As numerical surrogates, FE head models can resolve geometrical and mechanical complexities of the human head as well as interfacial conditions among various intracranial components. Many FE models have been established over the past decades in light of their immense potential to explore the pathomechanical cascades from external insults, localized tissue responses, and resultant brain injury (see review articles (Giudice et al., 2019; Madhukar and Ostoja-Starzewski, 2019; Yang and Mao, 2019)). Particularly, advances in neuroimaging, that delineates intracranial structures in a non-invasive fashion, largely accelerate the improvement of anatomical representations in contemporary head models. For example, by leveraging high-resolution computed tomography scans and angiography, several three-dimensional (3D) head models have successfully captured cranial morphology (De Kegel et al., 2019; Li et al., 2017, 2019) and cerebral vasculature (Ho and Kleiven, 2007; Subramaniam et al., 2021; Zhao and Ji, 2020), respectively. Such FE models with high-level structural details often possess fine meshes in order to appropriately resolve the geometrical complexities. For instance, two recent brain models with conforming hexahedral meshes that captured the morphological heterogeneity of the cerebral cortex (i.e., gyri and sulci) and lateral ventricles exhibited a mean mesh size of 0.85 mm with up to 1.2 million elements for the brain (Li et al., 2021; Zhou et al., 2022; Zhou et al., 2019b). For head models with smoothed brain surfaces, the number of brain elements varied from 5 k to 202 k with mesh resolutions of 1.8 – 6 mm (Chatelin et al., 2011; Kleiven and von Holst, 2002; Mao et al., 2013; Zhao and Ji, 2019a; Zhou et al., 2016).

Partially inspired by the experimental findings that functional impairment and morphological damage of axonal fibers were relevant to the strain regime imposed on the experimental tissue (LaPlaca et al., 2007; Montanino, 2020; Morrison III et al., 2011; Wu et al., 2021b), fiber orientation delineated by diffusion tensor imaging (DTI) has been integrated into FE models in recent years to inform the computation of axonal fiber deformation. Thus far, various techniques have been proposed for such an integration. For example, Zhou et al. (2021a) projected the strain tensor along the realtime fiber orientation to obtain the deformation along the fiber tracts (i.e., tract-oriented strain, which is alternatively termed as axonal strain or fiber strain in the literature), while Giordano et al. (2014) instead simulated the brain as a hyper-viscoelastic fiber-reinforced anisotropic medium with the fiber orientation integrated into the constitutive law to inform the fiber reinforcement. In these two referred studies, a critical issue of resolution mismatch between FE and DTI voxel exists. To address this, the voxel-wise fiber orientation in DTI was spatially downsampled and then implemented into FE models with one synthetic orientation per element (referred to as element-wise orientation implementation, hereafter). The process of spatially resampling orientation information in DTI voxels to match the FE resolution is referred to as orientation downsampling. This or similar element-wise orientation implementation technique has been used in other studies (Chatelin et al., 2011; Colgan et al., 2010; Giordano and Kleiven, 2014; Giordano et al., 2017; Hernandez et al., 2019; Kraft and Dagro, 2011; Kraft et al., 2012; Laksari et al., 2020; Sahoo et al., 2016; Sullivan et al., 2015; Zhao and Ji, 2019b; Zhou et al., 2021b). To circumvent the orientation downsampling, Garimella et al. (2019) explicitly incorporated the whole WM fiber tractography reconstructed from DTI into a head model in which multiple fibers were embedded within one WM element, while Ji et al. (2015) transformed the fiber orientation of WM voxels from the DTI space into the coordinate system of the FE head model then computed the tract-oriented strain at the voxel level by resolving the strain tensor of the WM element from pre-computed simulation to the fiber orientation of these transformed voxels. Regardless of these above-described disparities as well as other model-specific choices, maximum tract-oriented strain has been congruently reported as an appropriate measure of injury by several independent groups (Giordano and Kleiven, 2014; Wu et al., 2021a; Zhao et al., 2017; Zhou et al., 2021b). More recently, Zhou et al. (2021b) proposed three new tract-related strain metrics, measuring the normal deformation perpendicular to the fiber tracts (i.e., tract-perpendicular strain), and shear deformation along and perpendicular to the fiber tracts (i.e., axial-shear strain and lateral-shear strain, respectively), all of which exhibited superior injury predictability over maximum principal strain. Note that the tract-oriented strain, tract-perpendicular strain, axial-shear strain, and lateral-shear strain are collectively referred to as tract-related strains, each of which characterizes one aspect of the loading regime endured by the fiber tracts.

Given that the computation of any tract-related strain is directly dependent on the fiber orientation, appropriate integration of DTI-delineated fiber orientation into FE models is crucial. When limited to the element-wise orientation implementation involving orientation downsampling, the validity of the orientation downsampling operation has not been appropriately verified. For example, Giordano et al. (2014) averaged the fiber orientation obtained from DTI with a resolution of 2 mm × 2 mm × 3.6 mm to match the resolution of an FE model with an average element size of 6 mm. Similarly, Sahoo et al. (2014) downsampled the fiber orientation from a DTI atlas with an isotropic resolution of 1 mm^3^ for another FE model with mesh sizes ranging from 1.14 mm to 7.73 mm. Nevertheless, neither study has quantitatively compared the downsampled orientation information with its counterparts originally conveyed in DTI. Moreover, how the downsampling procedures affect the FE-derived tract-related strains remains to be systematically assessed.

Thus, the aim of the current study is to quantitatively evaluate the consequence of orientation downsampling on the computed WM tract-related deformation. To achieve this, a new approach was proposed to integrate the voxel-wise fiber orientation in DTI into FE models by embedding fiber orientations from multiple voxels within one element. This new approach and two existing downsampling algorithms were respectively implemented into two FE head models with different levels of mesh sizes. By setting the responses determined by the newly proposed voxel-wise implementation as the reference, the reliability of two alternative downsampling approaches was quantitatively evaluated. We thus tested the hypothesis that orientation downsampling causes a systematical loss of orientation information and further compromises the computation of tract-related strains.

## 2. Methods

### 2.1. Finite element brain models

Two previously developed head models with different mesh resolutions were used in this study, i.e., the KTH model (Kleiven, 2007) (**Fig. 1A-B**) and the ADAPT model (Li et al., 2021) (**Fig. 1C-D**). Each model consists of critical head components, including gray matter, WM, skull, scalp, subarachnoid cerebrospinal fluid (CSF), ventricles, superior sagittal sinus, transverse sinus, meninges, falx, and tentorium. The brain in both models was discretized by hexahedral elements with spatially heterogeneous element sizes. Following the tissue classification protocol proposed by Zhou et al. (2021b), the KTH model includes 1197 WM elements with a resolution of 6.37 ± 1.60 mm (range of 1.96 – 9.83 mm); the ADAPT model contains 113678 WM elements with a mesh size of 1.28 ± 0.55 mm (range of 0.34 – 3.23 mm). Numerical commonalities and differences between these two models are summarized in **Table A1** in Appendix as well as previous studies (Kleiven and von Holst, 2002; Li et al., 2021). Responses of both models have shown good correlation with experiments of brain strain, brain-skull relative motion, and intracranial pressure (Kleiven, 2006; Li et al., 2021; Zhou et al., 2019c; Zhou et al., 2019d; Zhou et al., 2018). Both models have been used to uncover the brain injury mechanism (Kleiven and von Holst, 2002; Li, 2021; Zhou et al., 2019a), evaluate the brain injury risk (Kleiven, 2007; Zhan et al., 2021), and inform the contrivance of advanced protective devices and treatment strategies (Fahlstedt et al., 2021; Wang et al., 2020).

**Fig. 1.**
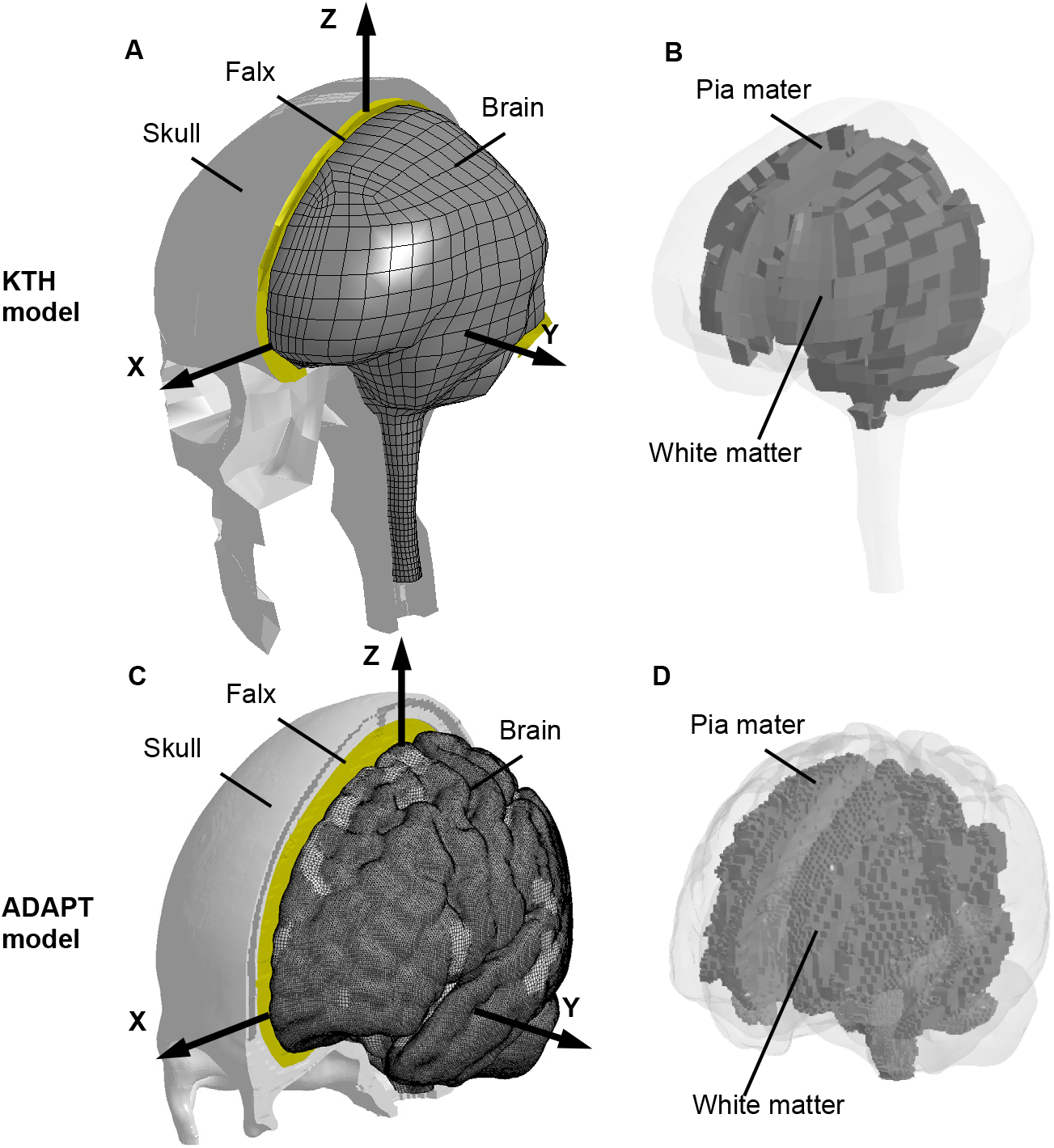
Two finite element head models with varying mesh resolutions. For the KTH head model with relatively coarse mesh, an isometric view of the head with the skull open to expose the brain is presented in subfigure (**A**) in which a skull-fixed coordinate system with the origin at the head’s center of gravity, while the pia mater and white matter are shown in subfigure (**B**) with the pia mater in translucency. Similar views are presented for the ADAPT head model with relatively fine mesh in subfigures (**C**) and (**D**). The brain FE elements are shown with mesh outlines in subfigures (**A**) and (**C**) to illustrate the mesh resolution.

### 2.2. Implementation of element-wise fiber orientation into FE brain models

To compute WM fiber tract-related deformation, fiber orientation from an open-access ICBM DTI-81 atlas (181 × 217 × 181 voxels, 1 mm isotropic, **Fig. 2A**) (Mori et al., 2008) was respectively integrated into the KTH model and ADAPT model using two previously proposed downsampling approaches. As exemplified by the KTH model in **Fig. 2**, the brain mesh was voxelized to obtain a reference volume (**Fig. 2B**), which was further aligned with the brain volume of the ICBM DTI-81 atlas via a diffeomorphic Demon registration (Li et al., 2021) (**Fig. 2C**). Then, for each WM element in the brain model, corresponding voxels in the DTI atlas enclosed by the given element were identified (**Fig. 2D**). Similarly, a registration operation was also conducted to spatially align the ICBM DTI-81 atlas with the brain mask of the ADAPT model.

**Fig. 2.**
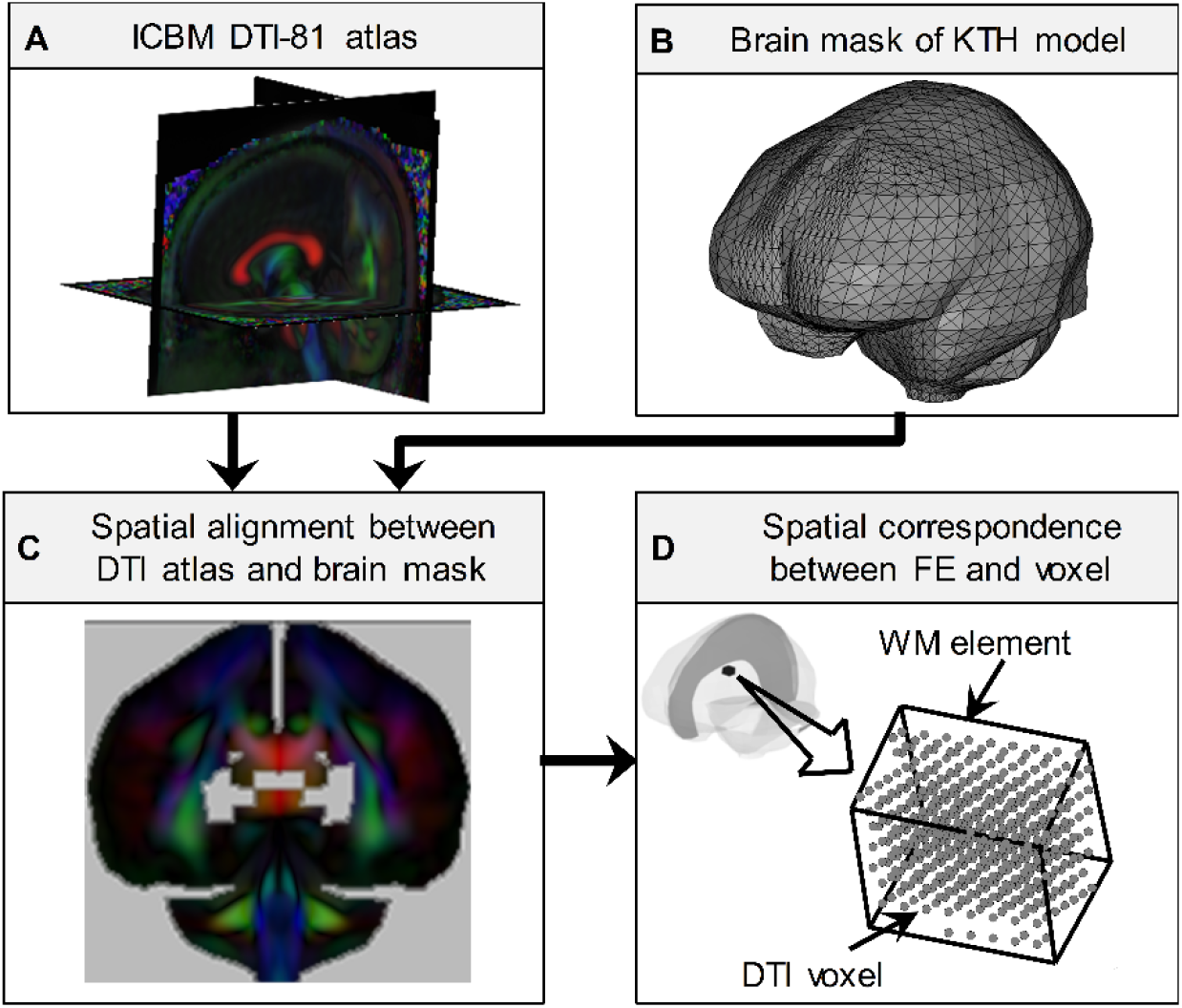
Spatial alignment between the DTI volume (i.e., ICBM DTI-81 atlas in subfigure (**A**)) and the brain mask of KTH model (**B**) with the outcome shown in subfigure **C**. For each WM element in the brain model, corresponding voxels in the DTI atlas enclosed by the given element were identified, which is illustrated by one representative element in subfigure **D**. Note that the DTI voxels in subfigure **D** are illustrated as spheres to show the voxel positions, while these spheres do not indicate these voxels are structurally isotropic.

To integrate the fiber orientation from multiple voxels into each WM element (**Fig. 2D**) toward element-wise orientation implementation, either the first eigenvectors of the diffusion tensors or the diffusion tensors themselves in these identified voxels were respectively downsampled using equations (1) (referred to as vector-averaged approach) and (2) (referred to as tensor-averaged approach) via a distance-weighted averaging scheme. In this scheme, the information from those voxels closer to the centroid of the element was more heavily weighted:

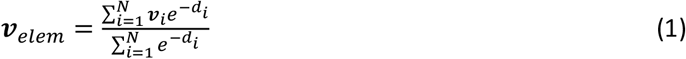

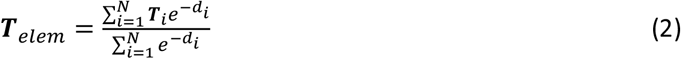

where ***ν**_elem_* and ***T**_elem_* are respectively the mean first eigenvector and the mean diffusion tensor calculated for each WM element; *N* is the number of selected voxels for the respective WM element; ***ν**_i_* and ***T**_i_* are respectively the first eigenvector and diffusion tensor of each selected voxel; *d_i_* is the distance from the voxel to the element centroid. Note that, before the implementation of equation (1), the angle between ***ν**_i_* and a preselected reference vector (i.e., the **X**-axis in the skull-fixed coordinate system in **Fig. 1** in the current study) was examined. Under the condition that the angle was greater than 90°, the direction of ***ν**_i_* would be flipped (i.e., ***ν**_i_*). As highlighted by Zhao and Ji (2019b), such a flip operation was needed since fiber orientation is expressed with either ***ν**_i_* or ***ν**_i_*, and it is necessary to ensure the consistency in the eigenvector across all the voxels and circumvent improper cancellation in equation (1).

The resultant orientation from equations (1) and (2) was respectively implemented at an element-wise basis. For the vector-averaged approach (equation 1), ***ν**_elem_* was considered the mean fiber orientation for each element (**Fig. 3A**), similar to the approach in Chatelin et al. (2011) and Zhao and Ji (2019b). For the tensor-weighted approach (equation 2), the first eigenvector of ***T**_elem_* was regarded as the mean orientation for each element (**Fig. 3B**), the same as the strategies in Zhou et al. (2021b) and Giordano et al. (2014).

**Fig. 3.**
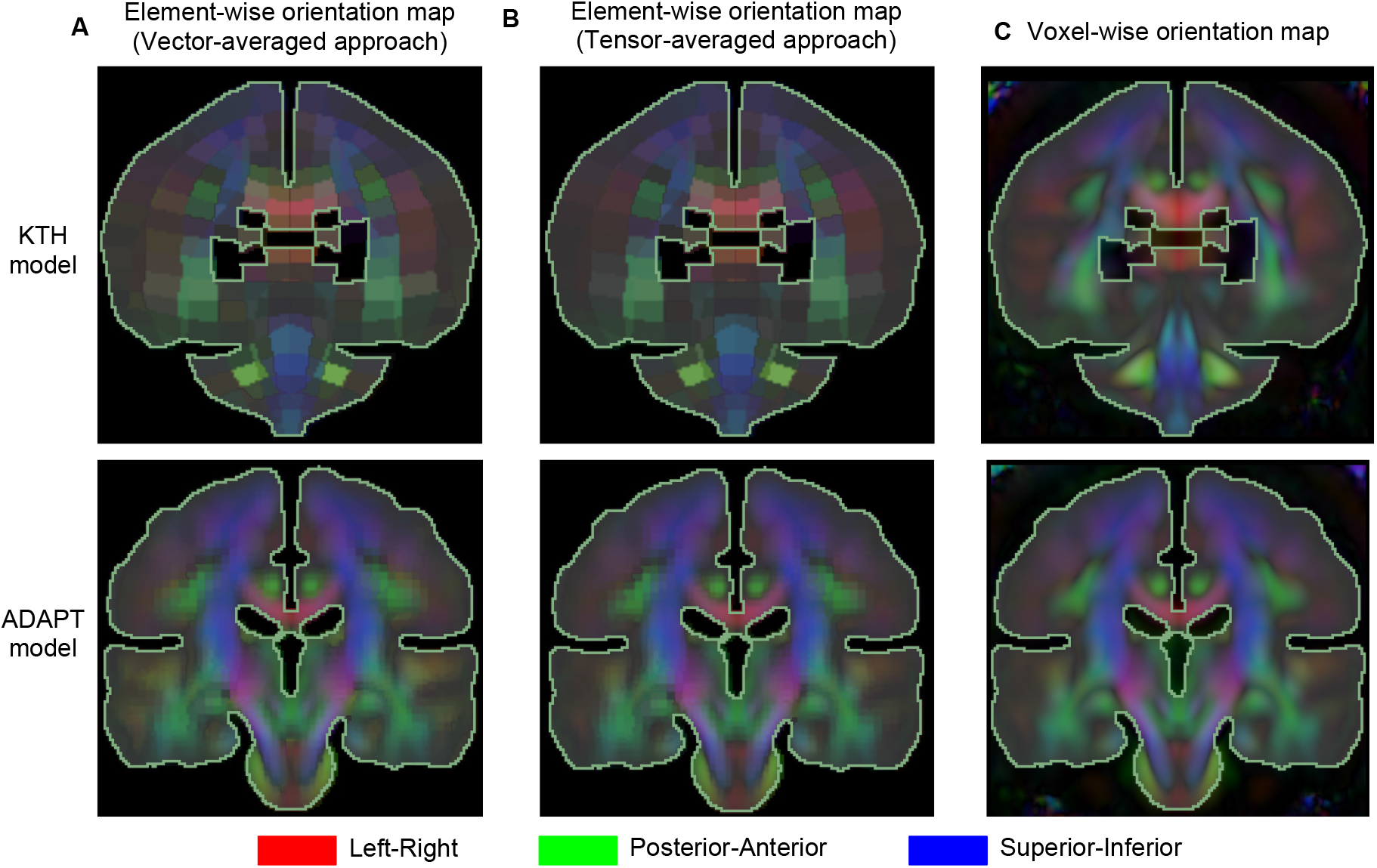
Color-coded orientation maps with the orientation implemented either at an element-wise basis using the vector-averaged approach (**A**) and tensor-averaged approach (**B**) or at a voxel-wise basis using the voxel-wise approach (**C**). The outline of the brain mask with the differentiation of ventricles used for image registration is superimposed to the orientation map.

### 2.3. Implementation of voxel-wise fiber orientation into FE brain models

To circumvent the downsampling procedures, we proposed a new approach by embedding orientations from multiple voxels within one element toward voxel-wise orientation implementation. In the same way as for the two downsampling approaches described above, DTI voxels enclosed by each WM element were identified at first based on their spatial correspondence (**Fig. 2D**). For each WM element, the first eigenvectors of all identified voxels (i.e., ***ν**_i_* in equation (1)) were wholly embedded into a given element without downsampling (see section 2.4). Thus, the fiber orientation was implemented at a voxel-wise basis (referred to as the voxel-wise approach hereafter) with an isotropic resolution of 1 mm^3^ given by the DTI atlas (**Fig. 3C**).

### 2.4. Real-time fiber orientation for the computation of tract-related strains

To inform the computation of tract-related strains with real-time fiber orientation, an embedded element approach was leveraged to track the real-time fiber orientation following the approach presented earlier (Zhou et al., 2021a). For all the three above-mentioned approaches, the orientation information was concretely represented by truss elements (serving as slave elements) which were embedded within the WM elements (serving as master elements). Note that the truss elements only served as instruments to monitor the temporal orientation of fiber tracts at each time step (Zhou et al., 2021a). They were simulated as a null constitutive model with nominal density and cross-sectional area (see **Table A2** in Appendix or a previous work (Zhou et al., 2021a)) and do not contribute any mechanical stiffness. Thus, the current study does not suffer from volume redundancy or stiffness redundancy, which are two commonly-seen drawbacks of the embedded element method (Garimella et al., 2019).

For both the vector-averaged and tensor-averaged approaches, only one truss element was embedded per WM element. In particular, for the vector-averaged approach, the truss element is oriented in the same direction as that of ***ν**_elem_*, while, for the tensor-averaged approach, the direction of the truss elements is aligned with the first eigenvector of ***T**_elem_*. For the voxel-wise approach, multiple truss elements were embedded in one WM element, in which truss elements angled along the direction of the first eigenvector of those voxels (i.e., ***ν**_i_* in equation (1)) within the given element. As is common across the three approaches, both ends of the truss elements fell within the boundaries of their master elements by adjusting the length of the embedded truss elements. Thus, nodal motion of a given truss element was governed exclusively by its master element. Consequently, the real-time fiber orientation during head impacts was reflected by the temporal direction of the truss element, which was updated at each solution cycle of the time-marching simulation.

To compute WM tract-related deformation, both the Green-Lagrange strain tensors for all WM elements and the orientation of embedded truss elements were iteratively obtained at each timestep from pre-computed simulations within the global coordinate system. The strain tensor of each WM element was further rotated to the coordinate systems with one axis aligned with the real-time fiber orientation, through which four strain-based metrics relating to the fiber deformation were extracted (**Fig. 3**), i.e., tract-oriented strain, tract-perpendicular strain, axial-shear strain, and lateral-shear strain. For the vector-averaged approach, the strain tensor of each WM element was only related to one realtime fiber orientation, which is also the case for the tensor-averaged approach. For the voxel-wise approach, the strain tensor of one given WM element was related to the temporal direction of all the truss elements encased by the given WM element.

The implementation of the embedded element method for tracking the real-time fiber orientation and the computation of the four tract-related strains are described with greater details in Zhou et al. (2021a) and Zhou et al. (2021b), respectively.

### 2.5. Impact simulation

To study the influence of fiber orientation downsampling on the computation of tract-related deformation, a concussive impact (i.e., case 157H2) from the National Football League dataset (Sanchez et al., 2019) was simulated using the KTH model and the ADAPT model with embedded fiber orientation implemented by the vector-averaged approach, tensor-averaged approach, and voxel-wise approach, respectively. In case157H2, the football player was laterally struck, resulting in high angular velocities in the coronal and sagittal planes. In the simulation, the translational and rotational velocities were prescribed to the rigid skull with the velocity profiles expressed with respect to the skull-fixed coordinate system. The impact was recorded for 50 ms with the computational time detailed in **Table 1**. The model responses were output at every 1 ms.

**Table 1.** Computational time for the KTH model and ADAPT model in combination with three orientation implementation approaches for a simulation of 50 ms solved by a massively parallel processing version of LS-DYNA with 128 CPUs.

| Time | Vector-averaged approach | Tensor-averaged approach | Voxel-wise approach |
| --- | --- | --- | --- |
| **KTH model** | 50 minutes | 50 minutes | 4 hours |
| **ADAPT model** | 33 hours | 33 hours | 67 hours |

### 2.6. Data analysis

Identical to those of the orientation information being integrated into the FE models, spatial resolutions of FE-derived responses were at the voxel level for the voxel-wise approach and at the element level for the vector- and tensor-averaged approaches. To address this disparity, all responses were mapped back to the imaging space of the ICBM DTI-81 atlas. This was intuitive for the voxel-wise approach, given that a one-to-one correspondence between the voxel and truss element was established. For the tensor- and vector-averaged approaches in which orientation information in multiple voxels was downsampled to be a single vector and further embedded in one WM element in the form of one truss element, responses based on the single truss element were assigned to all the voxels enclosed by the given WM element. Such mapping operations synchronized the response resolutions and facilitated direct comparisons among the three approaches. Response-encoded contours were generated in the imaging space.

To quantify the local fiber orientation embedded within the FE models, the skull-fixed coordinate system was selected as the reference. Adapting the strategy by Giordano et al. (2017), the fiber orientation was expressed in terms of two Eulerian angles, i.e., elevation angle (*α*) and azimuth angle (*β*) (**Fig. 5, Table 2A**). To identify the localized alteration of fiber orientation associated with the downsampling procedures by either the vector- or tensor-averaged approaches, absolute deviations in elevation angle and azimuth angle between the voxel-wise approach and each of the two downsampling approaches were calculated at the voxel-wise level, respectively (**Table 2B**).

**Table 2.**
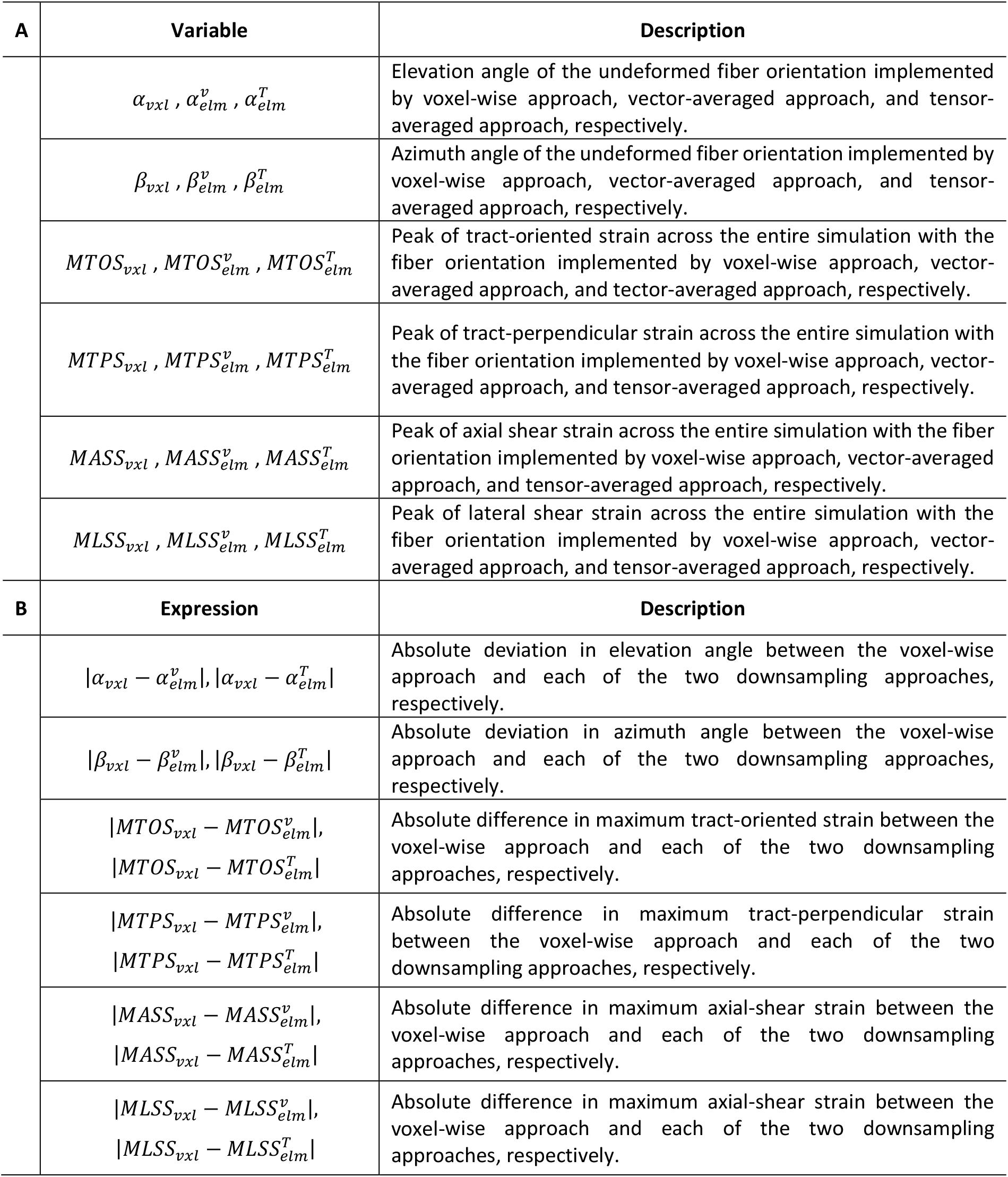
Summary of variables quantifying the fiber orientation implemented by three approaches and tract-related strains (**A**) and expression quantifying the absolute differences associated with the orientation downsampling (**B**). In each variable, the subscript indicates the basis that the orientation information is implemented (i.e., *vxl* for the voxel-wise basis, *elm* for the element-wise basis), while the superscript indicates the orientation downsampling approaches (i.e., *v* for vector-averaged approach, *T* for tensor-averaged approach).

To further evaluate the consequence of orientation downsampling on the computation of fiber deformation, four tract-related strains (**Fig. 4** and **Table 2A**) were extracted from FE models with the real-time fiber orientation described above. For each of the four strains, the absolute difference between the voxel-wise approach and two downsampled approaches was respectively calculated per voxel (**Table 2B**) and was further expressed in the form of normalized root-mean-square error (NRMSE; normalized by the mean value determined by the voxel-wise approach). To quantify the influence of the orientation downsampling on the peak strain responses, the 95^th^ percentile maximum values of four tract-related strains were extracted across the whole WM. All data analyses were performed in MATLAB (2016b; Mathworks, Natick, MA).

**Fig. 4.**
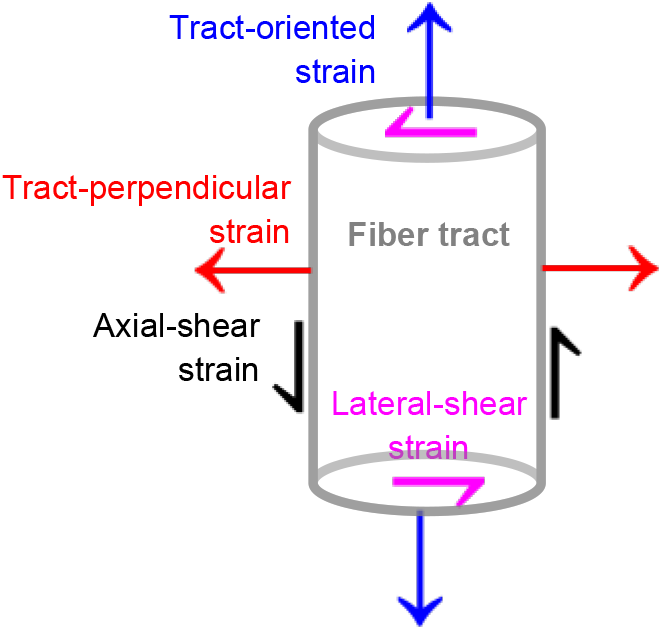
Illustration of the four tract-related strains with respect to the fiber tract. Tract-oriented strain and tractperpendicular strain measure the normal deformation along and perpendicular to the fiber tracts, while axial-shear strain and lateral-shear strain measure shear deformation along and perpendicular to the fiber tracts.

**Fig. 5.**
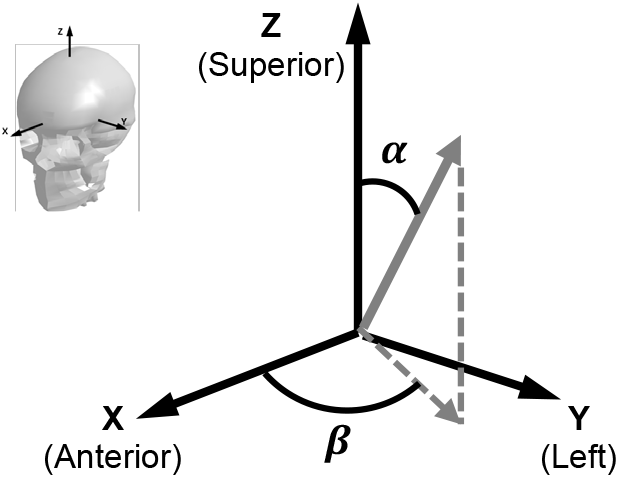
Characterization of the fiber orientation (gray arrow in solid line) by means of elevation angle (*α* ∈ [0°, 90°]) and azimuth angle *(β ∈* [0°, 90°]). These angles are expressed with respect to the skull-fixed coordinate system with a whole-head thumbnail on the top-left corner.

## 3. Results

To clarify the differences in elevation angle and azimuth angle quantifying the fiber orientation integrated into FE models and four tract-related strains measuring the deformation of fiber tracts among the three orientation implementation approaches, this section is organized in a hierarchical order to facilitate the readers’ understanding. We firstly illustrate the analysis based on the results of one randomly selected WM element of each FE model and then extend the analysis across the whole white matter region.

### 3.1 Effects of orientation downsampling on the responses of one representative element

For one representative WM element in the KTH model (**Fig. 6A-C**) encasing 361 voxels, the mean values for *α_vxl_* and *β_vxl_* in the voxel-wise approach were 64.1° (range of 47.8° – 89.5°) and 66.3° (range of 54.6° – 73.3°), respectively. Due to the downsampling operation, the resultant elevation and azimuth angles were 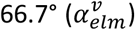 and 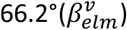 for the vector-averaged approach, and 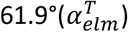 and 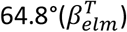 for the tensor-averaged approach, all of which fell within the range measured from the voxel-wise approach (**Fig. 6B**). Similarly, the range of each tract-related strain calculated from the voxel-wise approach covered its counterparts by the two downsampling approaches (**Fig. 6C**). Using the results determined by the voxel-wise approach as the reference, the NRMSE values ranged from 4.2% in MASS to 25.4% in MTPS for the vector-averaged approach, and from 4.7% in MASS to 25.9% in MTPS for the tensor-averaged approach.

**Fig. 6.**
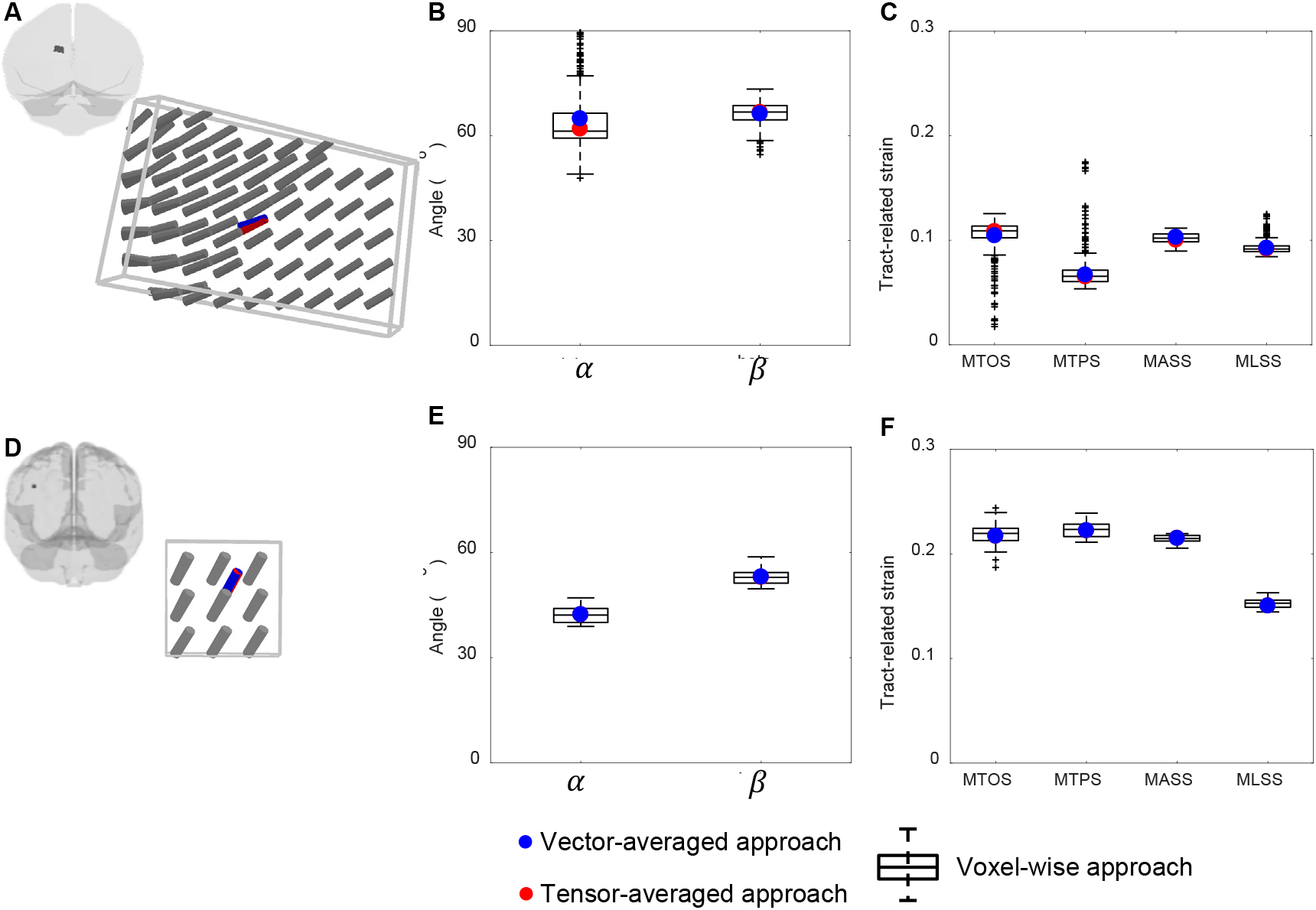
(**A**) Embedded truss elements implemented in one representative WM element in the KTH model with a whole-head thumbnail showing the view angle. The truss elements are coloured in blue and red for the vector- and tensor-averaged approaches, respectively, while in gray for the voxel-wise approach. (**B**) Boxplots of elevation angle (*α*) and azimuth angle (*β*) of the truss elements in subfigure (**A**). (**C**) Boxplots of four tract-related strains for the representative element in subfigure (**A**). Similar figures are also shown for one representative WM element in the ADAPT model in subfigures (**D-F**) in the right column. Note that in subfigures (**A**) and (**B**), the representative WM element enclosed 361 voxels for the KTH model and 32 voxels for the ADAPT model. Thus, some of the truss elements (in grey) implemented by the voxel-wise approach were overlapped with each other from the given view angles.

A similar analysis was performed on one randomly-selected element enclosing 36 voxels in the ADAPT head model (**Fig. 6D-F**). All the response variables of interest determined by the voxel-wise approach covered their counterparts from the two downsampling approaches. For fiber orientation, the voxel-wise approach (**Fig. 6D**) exhibited a range of 38.9° – 47.1° for *α_vxl_* (mean value: 42.3°) and 49.6° – 58.8° for *β_vxl_* (mean value: 53.1°), while fiber angles measured from the vector- and tensor-averaged approaches were commensurable with each other (*i.e*. 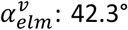 vs. 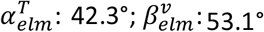 vs. 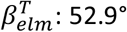) (**Fig. 6E**). For the strain-based metrics, the two downsampling approaches lead to nearly identical values in all four tract-related strains (**Fig. 6F**). The NRMSE values for the tensoraverage approach spanned from 1.8% (MASS) to 5.7% (MTOS) compared with the voxel-wise approach. The same NRMSE values were also noted for the vector-average approach when limiting the precision to one digit.

### 3.2. Effects of orientation downsampling on the responses of whole white matter region

The above-described analysis illustrated by one representative element per model was extended to the whole WM region. To quantify the extent of orientation downsampling in both brain models, the number of voxels identified for each WM element was examined. For the KTH model, the number of voxels per WM element was 312 ± 198 (range of 6-980) (**Fig. 7A**). Consequently, 1197 truss elements were respectively embedded per the implementation of vector- and tensor-averaged approaches, while 373976 truss elements for the voxel-wise approach (**Fig. 7C**). For the ADAPT model, a mean of 3 voxels (range of 1-57) was identified for a typical WM element (**Fig. 7B**). The numbers of truss elements introduced by both downsampling approaches were 113678, in comparison with 433299 by the voxel-wise approach (**Fig. 7D**).

**Fig. 7.**
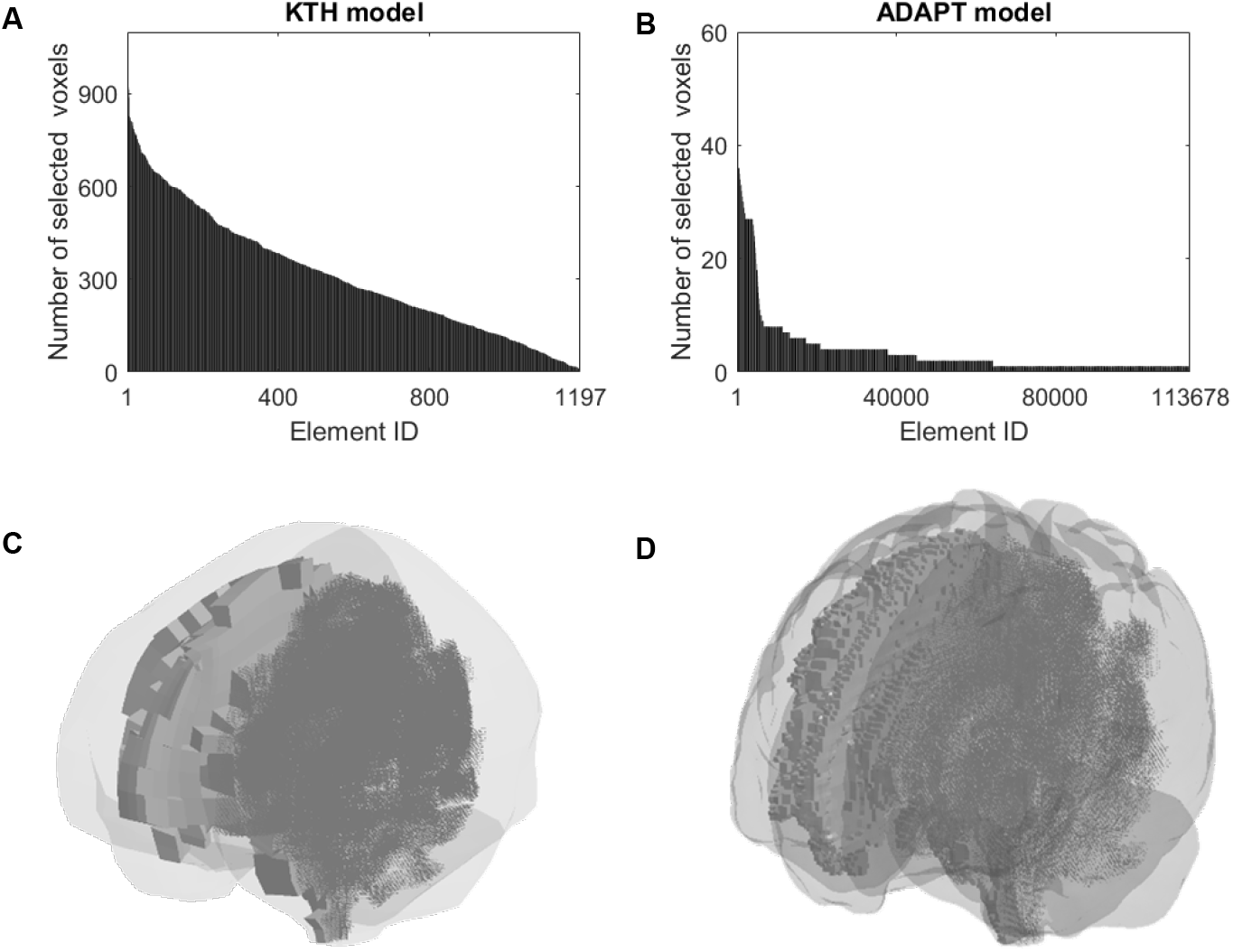
Number of DTI voxels selected for each of the WM element (**A-B**) and brain model with WM elements and embedded truss elements implemented by the voxel-wise approach (**C-D**) in the KTH model (left column) and ADAPT model (right column). For better illustration, the brain model in subfigures (**C**) and (**D**) is shown in translucency with the WM element in only one hemisphere being visible.

In both models, the three approaches resulted in similar distributions of *α* in the deep brain regions, such as the corpus callosum with the *α* consistently around 90°. Nevertheless, in the KTH model, the absolute deviation in *α* over 30° due to the orientation downsampling exhibited a patchy distribution, spreading across the WM/gray matter interface and the pons regions (highlighted in **Fig. 8A**). Such patchily-distributed differences were not seen in the ADAPT model with relatively fine meshes, in which only a handful of isolated voxels with absolute differences in *α* over 30° were revealed. Similar trends were also noted in *β* (**Fig. A1** in Appendix).

**Fig. 8.**
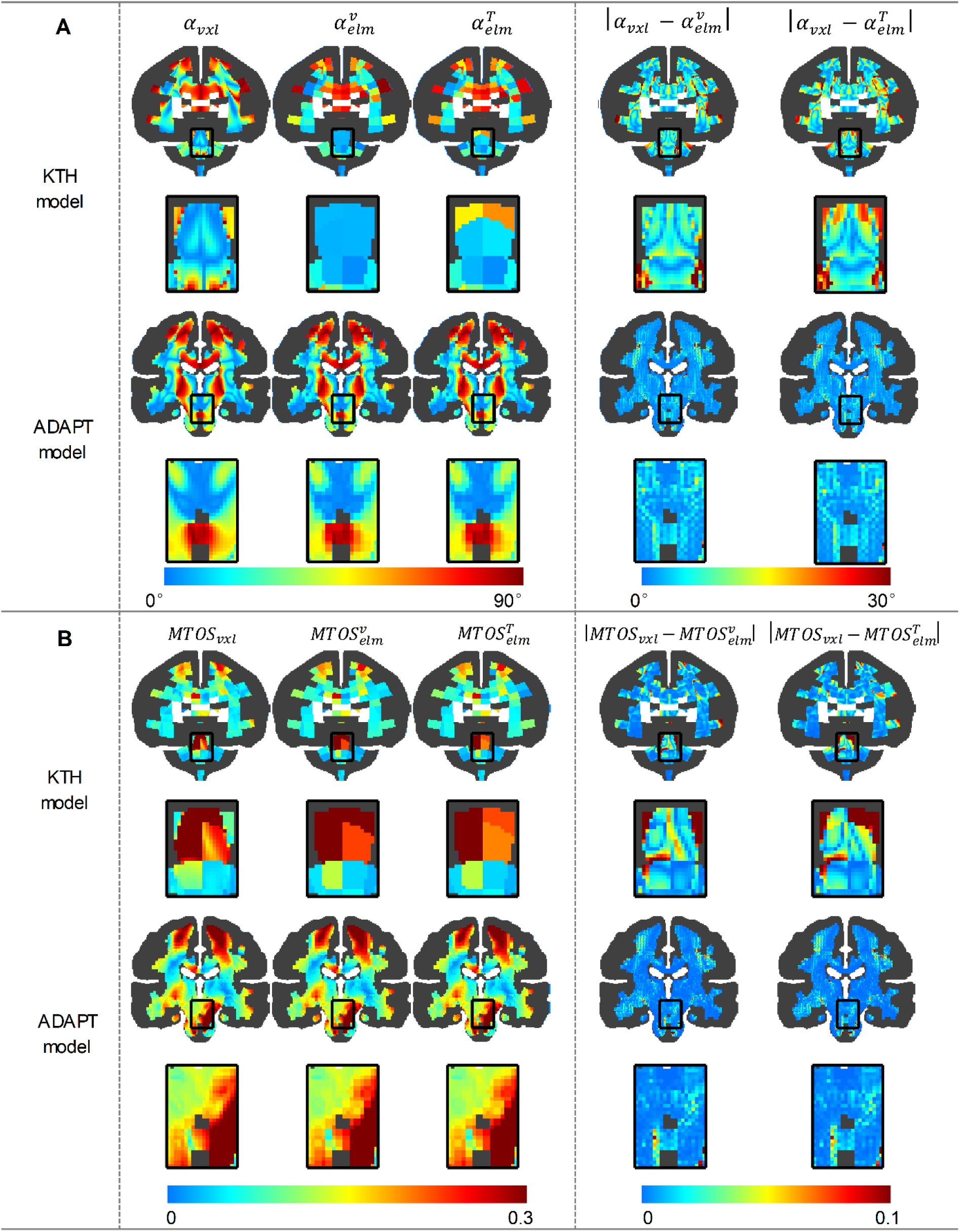
Coronal cross-sections of elevation angle (*α*) of fiber orientation (**A**) and MTOS (**B**) determined by the voxel-wise approach and two downsampling approaches along with the absolute difference associated with fiber orientation downsampling. Each cross-section is provided at the whole-brain level along with an enlarged view for the pons regions, in which the gray matter region is colored in dark gray.

To evaluate the consequence of orientation downsampling on the computation of fiber deformation, four tract-related strains were calculated from both models with the fiber orientation either directly implemented at the voxel-wise level or downsampled to the element-wise level. For the KTH model (**Fig. 8B**), both downsampling approaches introduced considerable differences (i.e., absolute difference exceeding 0.1) in MTOS, particularly in the pons regions. The NRMSE values across the WM region were 36.0% and 35.2% for the vector- and tensor-averaged approaches, respectively (**Table 3**). Similar trends were also noted for the other three tract-related strains (**Fig. A2** in Appendix). Quantitatively, the NRMSE values of other three tract-related strains across all WM region ranged from 19.0% (MLSS) to 22.8% (MTPS) for the vector-averaged approach, and from 18.5% (MLSS) to 22.0% (MTPS) for the tensor-averaged approach (**Table 3**).

**Table 3.** NRMSE in tract-related strains associated with fiber orientation downsampling in the KTH and ADAPT models. (**A**) Voxel-wise approach vs. Vector-averaged approach; (**B**) Voxel-wise approach vs. Tensor-averaged approach. The NRMSE values were computed across all WM voxels.

| **A** | NRMSE (%) | KTH model | ADAPT model | **B** | NRMSE (%) | KTH model | ADAPT model |
| --- | --- | --- | --- | --- | --- | --- | --- |
| Voxel-wise approach<br>vs.<br>Vector-averaged approach | *MTOS* | 36.0 | 4.5 | Voxel-wise approach<br>vs.<br>Tensor-averaged approach | *MTOS* | 35.2 | 4.6 |
|  | *MTPS* | 22.8 | 3.5 |  | *MTPS* | 22.0 | 3.5 |
|  | *MASS* | 20.6 | 3.2 |  | *MASS* | 20.1 | 3.1 |
|  | *MLSS* | 19.0 | 2.8 |  | *MLSS* | 18.5 | 2.8 |

When switching to the ADAPT model, orientation downsampling only introduced absolute differences of MTOS over 0.1 to a few voxels (**Fig. 8B**). This was further confirmed by the NRMSR value based on MTOS (**Table 3**), i.e., 4.5% for the vector-averaged approach and 4.6% for the tensor-averaged approach. Consistent with that of MTOS, the NRMSE values for the other three tract-related strains in the ADAPT model were consistently less than 5% both for the vector- and tensor-averaged approaches (**Table 3**). Strain contours of the resting three tract-related strains for the ADATP model are shown in **Fig. A2** in Appendix.

We further quantified the influence of the orientation downsampling on the peak tract-related deformation by extracting the 95^th^ percentile value of maximum tract-related strains across the whole WM region (**Fig. 9**). When limiting the comparison to the two downsampling approaches, only minor differences (less than 1%) were found between them independent of the strain types and FE models.

**Fig. 9.**
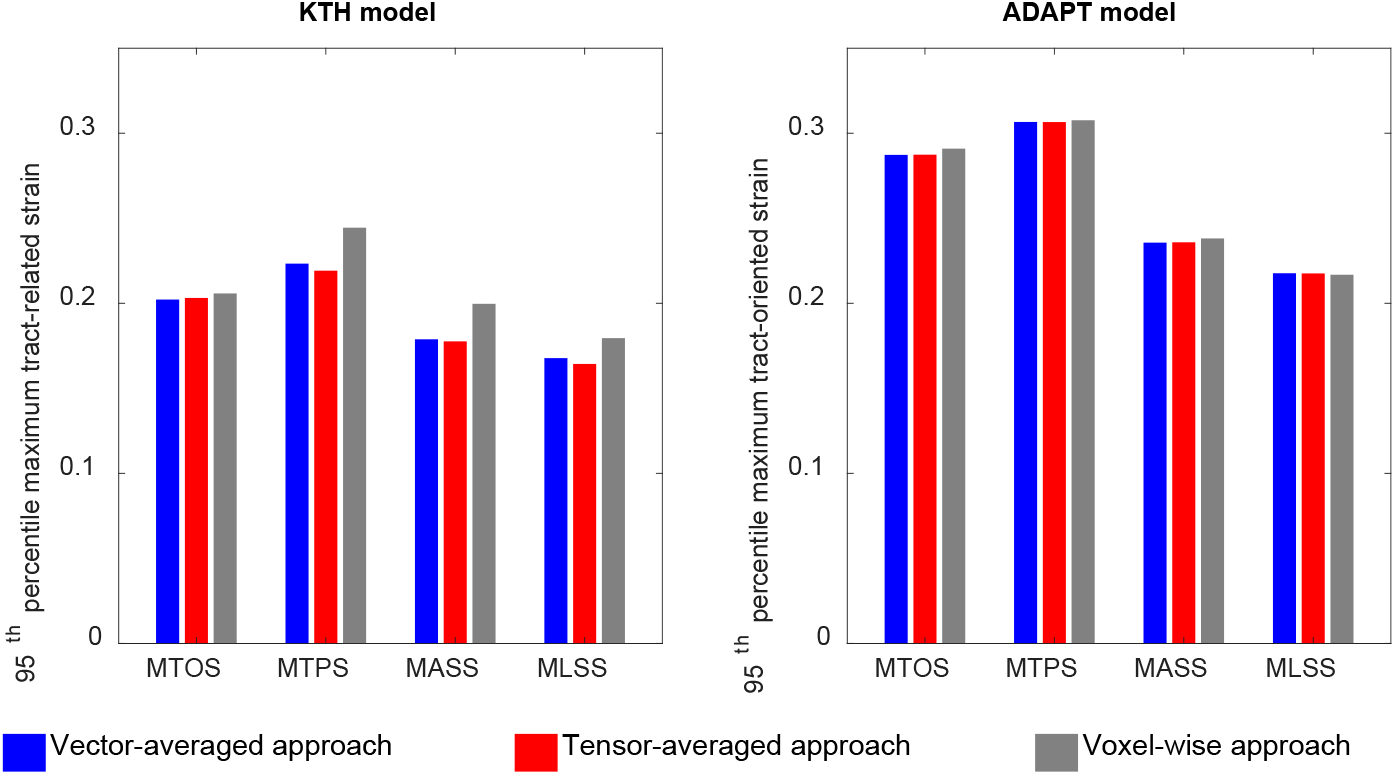
Comparison of 95^th^ percentile maximum tract-related strains determined by the voxel-wise approach and the two downsampling approaches in the KTH and ADAPT models.

When extending the comparison by incorporating the voxel-wise approach, the tensor-averaged downsampling approach underestimated the peak strains in the KTH model ranging from 1.7% in MTOS to 12.4% in MASS. For the ADAPT model, the differences in the peak values between the voxel-wise approach and the two downsampling approaches were consistently less than 1%.

## 4. Discussions

The current work proposed a new approach to incorporate voxel-wise fiber orientation from DTI into FE models by embedding orientations from multiple voxels within one element and evaluated the reliability of two previously-used orientation downsampling approaches with one orientation per element. By setting the responses from the voxel-wise implementation as the reference, it was verified that, for a model with a large mesh-image resolution mismatch, the orientation downsampling introduced an absolute deviation of fiber orientation over 30° across the WM/gray matter interface and the pons regions. This further compromised the computation of tract-related strains with the NRSMR values up to 30% and underestimated the peak tract-related strains up to 10%. For the other FE model with finer meshes, the downsampling-induced effect was lower, both on the fiber orientation and tract-related strains. Taken together, the voxel-wise orientation implementation is recommended in future studies as it leverages the DTI-delineated fiber orientation to a larger extent than the element-wise orientation implementation.

### 4.1 Implementing DTI-delineated fiber orientation into finite element brain models

In light of their orientation-dependent natures, tract-related strains require fiber orientation information to inform their computation. Although DTI has been available for decades to delineate axonal tracts, challenges remain on how to adequately integrate the high-resolution fiber orientation into FE models with nonidentical mesh resolutions. In preference to the element-wise orientation implementation with only one fiber orientation embedded within one WM element (e.g., the vector- and tensor-averaged approaches evaluated in the current work), the voxel-wise approach proposed in the current study addressed the mesh-imaging resolution mismatch by embedding orientation from multiple voxels within one element. This voxel-wise approach further served as the reference to evaluate the reliability of the vector- and tensor-averaged approaches. As enumerated by the NRMSE values and underestimated peak strain responses, our work quantified the loss of orientation information due to downsampling and the consequently compromised computation of tract-related strains, which is particularly true for the model with coarse meshes.

When comparing the results across all three approaches (one with, and two without downsampling), it was found that responses of these two downsampling approaches fell within the range of their counterparts by the voxel-wise approach, as partially demonstrated by one representative element per model (**Fig. 6**). Such findings provided certain support to the vector- and tensor-averaged approaches and indicated that the results in previous studies based on downsampled approaches were not absurdly inaccurate, such as the one by Zhou et al. (2021a) using the tensor-averaged approach and the other by Chatelin et al. (2011) using the vector-averaged approach. Despite this, incorporating voxel-wise fiber directions without downsampling (e.g., the voxel-wise approach in the current study) provides a better representation of WM fiber orientation and should be regarded as the rule of thumb.

When limiting the comparison to the results revealed by the vector- and tensor-averaged approaches, these two schemes performed alike, regardless of the FE models. This was substantiated by the similar distribution and magnitude of orientation information and tract-related strains revealed by these two approaches (**Fig. 8**, **Fig. 9**, and **Table 3**). Such similar performances were expected given that both approaches practically downsampled the same orientation information from DTI, with the only difference being the sequential order of eigenvector computation and the distance-weighted averaging. For the vector-averaged approach, the principal eigenvectors of tensors in these voxels enclosed by one WM element were computed at first. Then all eigenvectors were averaged by a set of weighting factors based on the distances between the DTI voxels and elemental centroid, outputting a single synthetic vector as the mean fiber orientation of the given element. For the tensor-averaged approach, the tensors of all enclosed DTI voxels were averaged at first with the output as a synthetic tensor, and then the principal eigenvector of the synthetic tensor was regarded as the mean fiber direction. In the current study, the same weighting factors were used in both averaging approaches, which further explained the similar performances between the vector- and tensor-averaged approaches.

Compared with the voxel-wise approach proposed in the current study, another promising alternative that could largely leverage the DTI-delineated orientation information is to embed the whole fiber tractography with continuous trajectories into FE models (Garimella, 2017; Hajiaghamemar and Margulies, 2021; Hajiaghamemar et al., 2020; Wu et al., 2019; Zhao et al., 2016). Despite its great potential, this approach conforms to multiple challenges associated with tractography reconstruction (Maier-Hein et al., 2017). Thus, the voxel-wise approach proposed in the current study can be regarded as an alternative to the previous tractography-based efforts, collectively reinforcing further the importance of leveraging the maximum amount of orientation information into the FE models toward an authentic prediction of tract-related deformation.

### 4.2 Limitations and path forward

Although this study yielded new insights on how to integrate the neuroimage-delineated fiber orientation into FE models, certain limitations should be acknowledged. First, besides the vector- and tensor-averaged approaches evaluated in the current study, other orientation downsampling strategies exist in the literature. For example, Li et al. (2021) implemented the principal eigenvector from the voxel closest to the centroid of the element as the mean fiber direction for the given element, while Kraft and Dagro (2011) added the principal eigenvectors of all voxels encased by the element with the outcome as the fiber direction. It is expected that these two referred approaches would perform similarly to the vector- and tensor-averaged approaches. Secondly, the current study simulated the brain as an isotropic and homogeneous medium, as no complete consensus has been reached yet about the mechanical heterogeneity and anisotropy of the human brain tissue. Relevant limitations have been exhaustively discussed in our recent publications (Zhou et al., 2021a; Zhou et al., 2021b) and are not repeated here. Nevertheless, we acknowledge that extending the current study to the context of modelling the brain as an anisotropic and heterogeneous object can be another interesting aspect for future work with the effort initiated by Zhao and Ji (2019b). Thirdly, across all the three approaches investigated in the current study, the orientation information of WM fiber tracts is based on the primary eigenvector of a rank-2 symmetric tensor. It has been well recognized that the tensor model is a limited approximation of fiber architecture, especially in those regions with fiber bundles intertwined and crossed (Jeurissen et al., 2013). The current study can be extended by exploiting more advanced models (Frank, 2001; Tournier et al., 2004) that better describe the diffusion in voxels with multidirectional fiber tracts. Lastly, although the voxel-wise approach proposed in the current study addresses the resolution mismatch between DTI voxel and FE, this approach remains to confront another resolution mismatch between strain tensors of brain elements (element-wise basis) and fiber orientation reflected by the direction of embedded truss elements (voxel-wise basis). This is reflected by the fact that the strain tensor of one given WM element was related to the temporal direction of all the truss elements encased by the given WM element to compute tract-related strain. Future work is planned to address this critical aspect with one alternative solution recently published by Ji and Zhao (2022).

## 5. Conclusion

The present study proposed a new approach to incorporate voxel-wise fiber directions from DTI into FE models by embedding fiber orientations from multiple voxels within one element and evaluated the reliability of orientation downsampling with one fiber orientation per element. This newly proposed approach and two existing downsampling alternatives were respectively implemented into two FE models with varying levels of element sizes. Two orientation downsampling approaches were shown to cause systematical loss of fiber orientation information and further compromised the accuracy of tract-related strain, particularly for the model with a large mesh-imaging resolution mismatch. For the model with refined meshes, these downsampling-induced effects were partially compensated. Collectively, the voxel-wise orientation implementation provides a better representation of fiber orientation and is recommended to be used in the future. Therefore, this study yields insights on integrating neuroimaging-informed fiber information into FE models to improve the accuracy of FE-derived WM tract-related deformation.

## Acknowledgments

This research has received funding from KTH Royal Institute of Technology (Stockholm, Sweden), the China Scholarship Council, the Swedish Foundation for International Cooperation in Research and Higher Education (STINT), and the Swedish Research Council (VR-2020-04496 and VR-2020-04724). The content of this article is solely the responsibility of the authors and does not necessarily represent the official views of funding agencies. The simulation were enabled by resources in project [SNIC 2021/5-406] provided by the Swedish National Infrastructure for Computing (SNIC) at the center for High Performance Computing (PDC), partially funded by the Swedish Research Council through grant agreement no. VR-2020-04496. The authors also thank for the anonymous reviewers for the stimulating comments and valuable suggestions that substantially improved this paper.

## Data availability

Scripts for mapping the fiber orientation information in diffusion tensor imaging to finite element brain models can be obtained by emailing request to the corresponding author.

## Author contributions

**Zhou Zhou:** Conception and study design, Finite element simulation, Imaging analysis, Data processing, Visualization; Writing – review & editing; **Teng Wang:** Conception and study design, Imaging analysis, Review & editing; **Daniel Jörgens:** Revision, Data analysis and illustration, Review & editing; **Xiaogai Li:** Conception and study design, Imaging analysis, Review & editing.

## Appendix

**Table A1.**
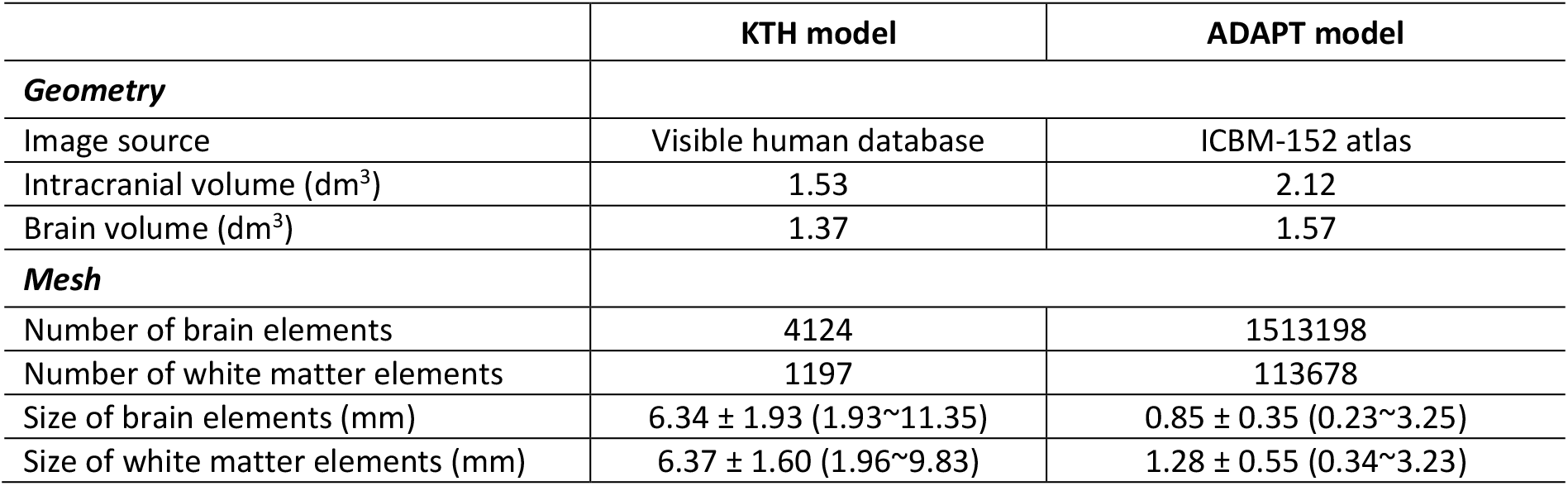
Summary of differences and commonalities between the KTH model and ADAPT model. Note that the mesh sizes are reported in the form of mean ± standard deviation (range).

**Table A2.**
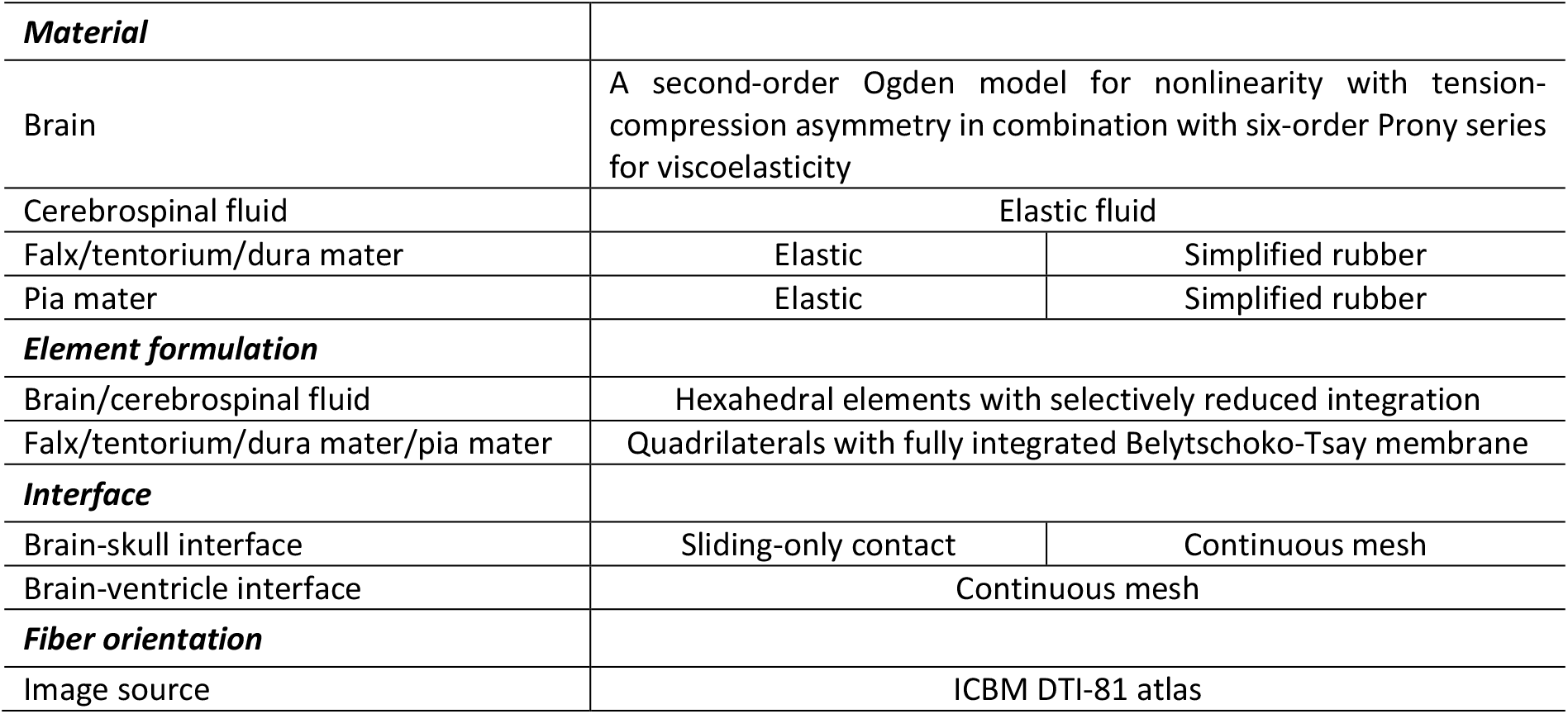
Material properties for the embedded truss elements.

**Fig. S1.**
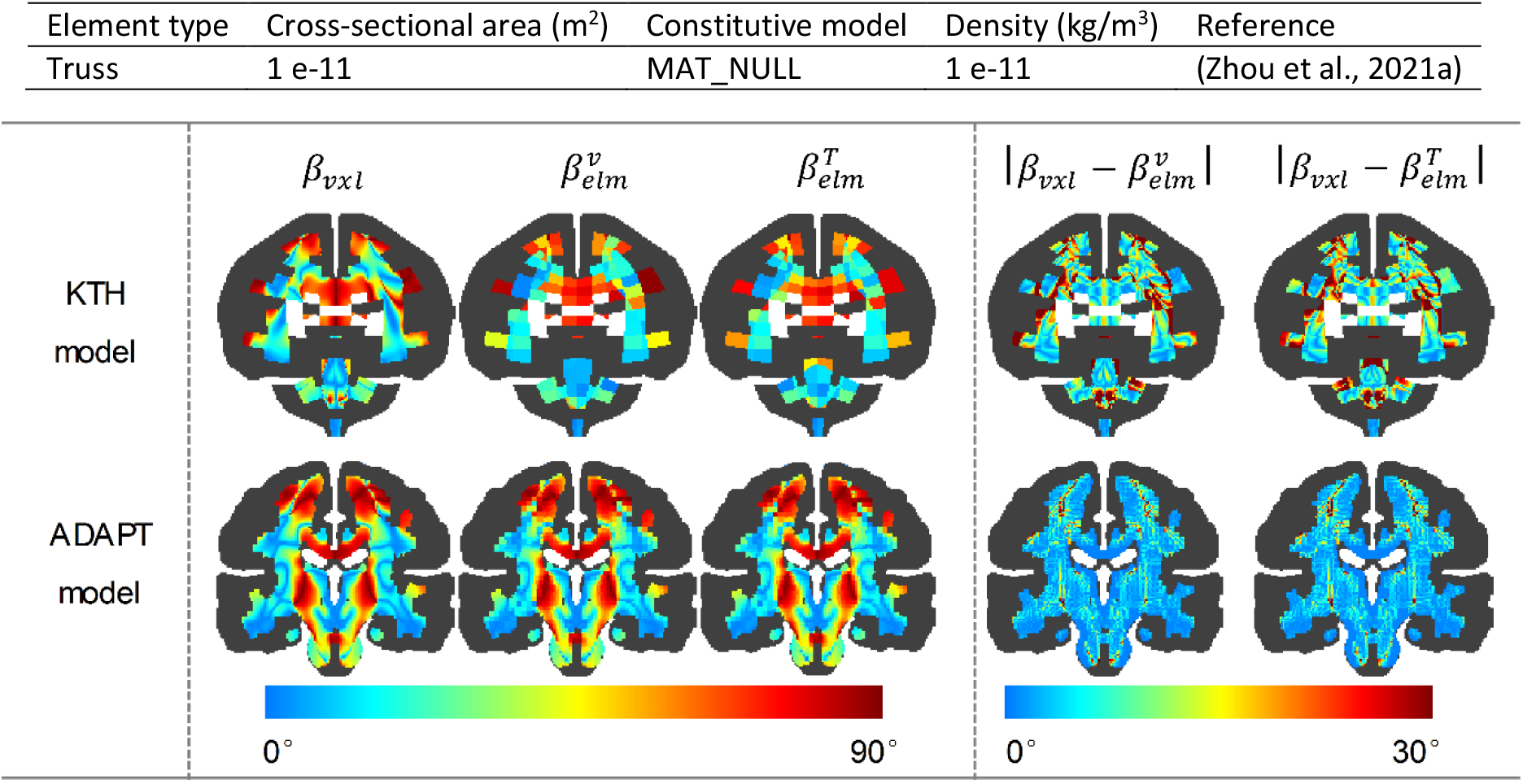
Coronal cross-sections of azimuth angle (*β*) of fiber orientation determined by the voxel-wise approach and two downsampling approaches along with the absolute difference associated with orientation downsampling. In each contour, the gray matter region is colored in dark gray.

**Fig. A2.**
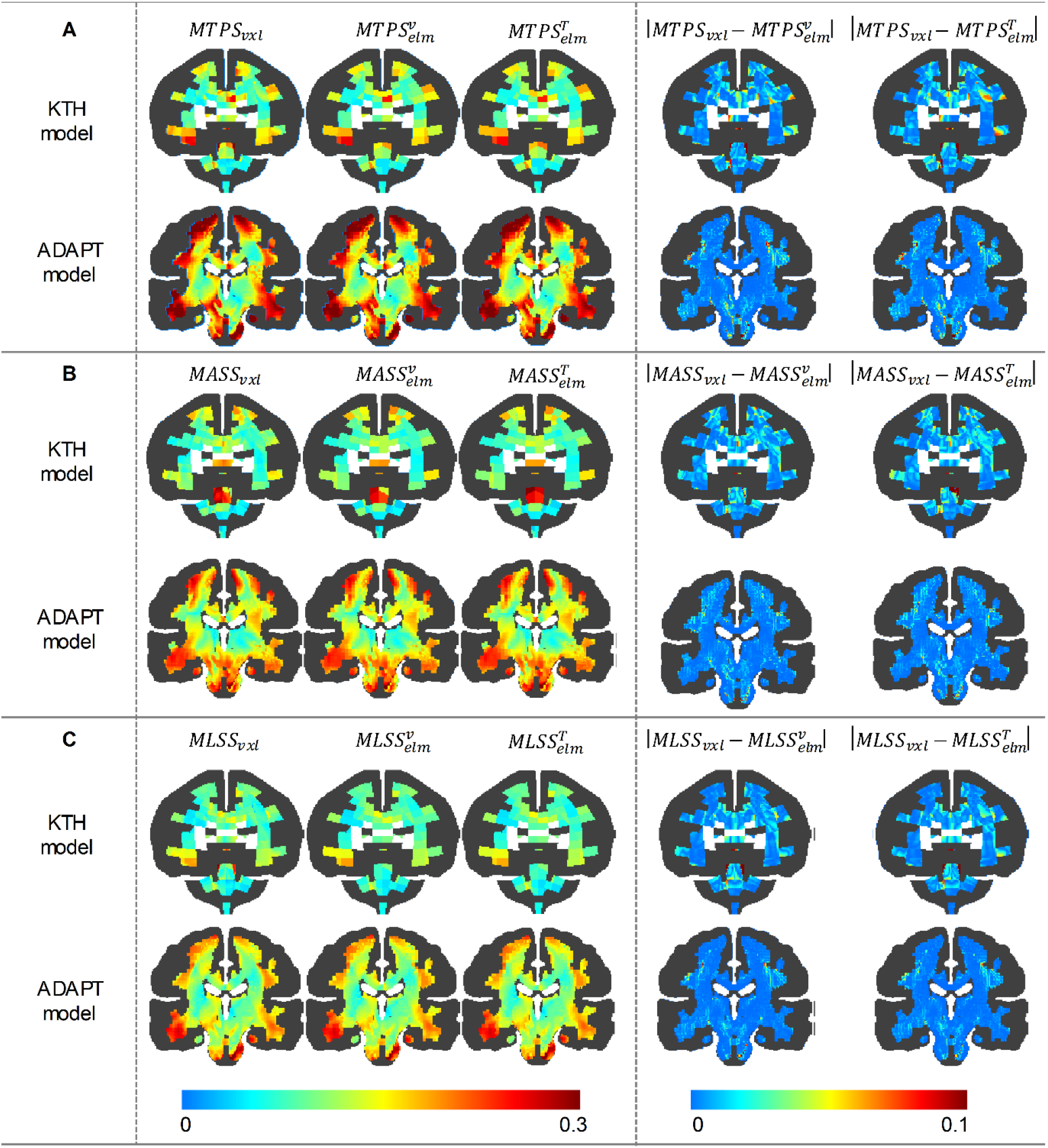
Coronal cross-sections of MTPS (**A**), MASS (**B**), and MLSS (**C**) determined by the voxel-wise approach and two downsampling approaches along with the absolute difference associated with orientation downsampling. In each contour, the gray matter region is colored in dark gray.

## References

Brazinova, A., Rehorcikova, V., Taylor, M.S., Buckova, V., Majdan, M., Psota, M., Peeters, W., Feigin, V., Theadom, A., Holkovic, L., 2021. Epidemiology of traumatic brain injury in Europe: a living systematic review. Journal of neurotrauma 38, 1411–1440.

Chatelin, S., Deck, C., Renard, F., Kremer, S., Heinrich, C., Armspach, J.-P., Willinger, R., 2011. Computation of axonal elongation in head trauma finite element simulation. Journal of the mechanical behavior of biomedical materials 4, 1905–1919.

Colgan, N.C., Gilchrist, M.D., Curran, K.M., 2010. Applying DTI white matter orientations to finite element head models to examine diffuse TBI under high rotational accelerations. Progress in biophysics and molecular biology 103, 304–309.

De Kegel, D., Meynen, A., Famaey, N., van Lenthe, G.H., Depreitere, B., Vander Sloten, J., 2019. Skull fracture prediction through subject-specific finite element modelling is highly sensitive to model parameters. Journal of the mechanical behavior of biomedical materials 100, 103384.

Fahlstedt, M., Abayazid, F., Panzer, M.B., Trotta, A., Zhao, W., Ghajari, M., Gilchrist, M.D., Ji, S., Kleiven, S., Li, X., 2021. Ranking and rating bicycle helmet safety performance in oblique impacts using eight different brain injury models. Annals of biomedical engineering 49, 1097–1109.

Frank, L.R., 2001. Anisotropy in high angular resolution diffusion – weighted MRI. Magnetic Resonance in Medicine: An Official Journal of the International Society for Magnetic Resonance in Medicine 45, 935–939.

Garimella, H.T., Menghani, R.R., Gerber, J.I., Sridhar, S., Kraft, R.H., 2019. Embedded finite elements for modeling axonal injury. Annals of biomedical engineering 47, 1889–1907.

Garimella, V.R.S.H.T., 2017. An embedded element based human head model to investigate axonal injury. The Pennsylvania State University.

Giordano, C., Cloots, R., Van Dommelen, J., Kleiven, S., 2014. The influence of anisotropy on brain injury prediction. Journal of biomechanics 47, 1052–1059.

Giordano, C., Kleiven, S., 2014. Evaluation of axonal strain as a predictor for mild traumatic brain injuries using finite element modeling. Stapp Car Crash J 58, 29–61.

Giordano, C., Zappalà, S., Kleiven, S., 2017. Anisotropic finite element models for brain injury prediction: the sensitivity of axonal strain to white matter tract inter-subject variability. Biomechanics and modeling in mechanobiology 16, 1269–1293.

Giudice, J.S., Zeng, W., Wu, T., Alshareef, A., Shedd, D.F., Panzer, M.B., 2019. An analytical review of the numerical methods used for finite element modeling of traumatic brain injury. Annals of biomedical engineering 47, 1855–1872.

Hajiaghamemar, M., Margulies, S.S., 2021. Multi-scale white matter tract embedded brain finite element model predicts the location of traumatic diffuse axonal injury. Journal of Neurotrauma 38, 144–157.

Hajiaghamemar, M., Wu, T., Panzer, M.B., Margulies, S.S., 2020. Embedded axonal fiber tracts improve finite element model predictions of traumatic brain injury. Biomechanics and modeling in mechanobiology 19, 1109–1130.

Hernandez, F., Giordano, C., Goubran, M., Parivash, S., Grant, G., Zeineh, M., Camarillo, D., 2019. Lateral impacts correlate with falx cerebri displacement and corpus callosum trauma in sports-related concussions. Biomechanics and modeling in mechanobiology 18, 631–649.

Ho, J., Kleiven, S., 2007. Dynamic response of the brain with vasculature: a three-dimensional computational study. Journal of biomechanics 40, 3006–3012.

Jeurissen, B., Leemans, A., Tournier, J.D., Jones, D.K., Sijbers, J., 2013. Investigating the prevalence of complex fiber configurations in white matter tissue with diffusion magnetic resonance imaging. Human brain mapping 34, 2747–2766.

Ji, S., Zhao, W., 2022. Displacement voxelization to resolve mesh-image mismatch: Application in deriving dense white matter fiber strains. Computer Methods and Programs in Biomedicine 213, 106528.

Ji, S., Zhao, W., Ford, J.C., Beckwith, J.G., Bolander, R.P., Greenwald, R.M., Flashman, L.A., Paulsen, K.D., McAllister, T.W., 2015. Group-wise evaluation and comparison of white matter fiber strain and maximum principal strain in sports-related concussion. Journal of neurotrauma 32, 441–454.

Kleiven, S., 2006. Evaluation of head injury criteria using a finite element model validated against experiments on localized brain motion, intracerebral acceleration, and intracranial pressure. International Journal of Crashworthiness 11, 65–79.

Kleiven, S., 2007. Predictors for traumatic brain injuries evaluated through accident reconstructions. Stapp car crash J 51, 81–114.

Kleiven, S., von Holst, H., 2002. Consequences of head size following trauma to the human head. Journal of biomechanics 35, 153–160.

Kraft, R.H., Dagro, A.M., 2011. Design and implementation of a numerical technique to inform anisotropic hyperelastic finite element models using diffusion-weighted imaging. ARMY RESEARCH LAB ABERDEEN PROVING GROUND MD WEAPONS AND MATERIALS RESEARCH ….

Kraft, R.H., Mckee, P.J., Dagro, A.M., Grafton, S.T., 2012. Combining the finite element method with structural connectome-based analysis for modeling neurotrauma: connectome neurotrauma mechanics.

Laksari, K., Fanton, M., Wu, L.C., Nguyen, T.H., Kurt, M., Giordano, C., Kelly, E., O’Keeffe, E., Wallace, E., Doherty, C., 2020. Multi-directional dynamic model for traumatic brain injury detection. Journal of neurotrauma 37, 982–993.

LaPlaca, M.C., Simon, C., Prado, G.R., Cullen, D., 2007. CNS injury biomechanics and experimental models. Progress in brain research 161, 13–26.

Li, X., 2021. Subject-specific head model generation by mesh morphing: A personalization framework and its applications. Frontiers in bioengineering and biotechnology.

Li, X., Sandler, H., Kleiven, S., 2017. The importance of nonlinear tissue modelling in finite element simulations of infant head impacts. Biomechanics and modeling in mechanobiology 16, 823–840.

Li, X., Sandler, H., Kleiven, S., 2019. Infant skull fractures: Accident or abuse?: Evidences from biomechanical analysis using finite element head models. Forensic science international 294, 173–182.

Li, X., Zhou, Z., Kleiven, S., 2021. An anatomically detailed and personalizable head injury model: Significance of brain and white matter tract morphological variability on strain. Biomechanics and modeling in mechanobiology 20, 403–431.

Madhukar, A., Ostoja-Starzewski, M., 2019. Finite element methods in human head impact simulations: a review. Annals of biomedical engineering 47, 1832–1854.

Maier-Hein, K.H., Neher, P.F., Houde, J.-C., Côté, M.-A., Garyfallidis, E., Zhong, J., Chamberland, M., Yeh, F.-C., Lin, Y.-C., Ji, Q., 2017. The challenge of mapping the human connectome based on diffusion tractography. Nature communications 8, 1–13.

Mao, H., Gao, H., Cao, L., Genthikatti, V.V., Yang, K.H., 2013. Development of high-quality hexahedral human brain meshes using feature-based multi-block approach. Computer methods in biomechanics and biomedical engineering 16, 271–279.

Maxwell, W., Dhillon, K., Harper, L., Espin, J., MacIntosh, T., Smith, D., Graham, D., 2003. There is differential loss of pyramidal cells from the human hippocampus with survival after blunt head injury. Journal of Neuropathology & Experimental Neurology 62, 272–279.

Menon, D.K., Schwab, K., Wright, D.W., Maas, A.I., 2010. Position statement: definition of traumatic brain injury. Archives of physical medicine and rehabilitation 91, 1637–1640.

Montanino, A., 2020. Definition of axonal injury tolerances across scales: A computational multiscale approach. Kungliga Tekniska högskolan.

Mori, S., Oishi, K., Jiang, H., Jiang, L., Li, X., Akhter, K., Hua, K., Faria, A.V., Mahmood, A., Woods, R., 2008. Stereotaxic white matter atlas based on diffusion tensor imaging in an ICBM template. Neuroimage 40, 570–582.

Morrison III, B., Elkin, B.S., Dollé, J.-P., Yarmush, M.L., 2011. In vitro models of traumatic brain injury. Annual review of biomedical engineering 13, 91–126.

Peterson, A.B., Xu, L., Daugherty, J., Breiding, M.J., 2019. Surveillance report of traumatic brain injury-related emergency department visits, hospitalizations, and deaths, United States, 2014. Centers for Disease Control and Prevention, U.S. Department of Health and Human Services, Atlanta, GA, pp. 1–23.

Sahoo, D., Deck, C., Willinger, R., 2014. Development and validation of an advanced anisotropic visco-hyperelastic human brain FE model. Journal of the mechanical behavior of biomedical materials 33, 24–42.

Sahoo, D., Deck, C., Willinger, R., 2016. Brain injury tolerance limit based on computation of axonal strain. Accident Analysis & Prevention 92, 53–70.

Sanchez, E.J., Gabler, L.F., Good, A.B., Funk, J.R., Crandall, J.R., Panzer, M.B., 2019. A reanalysis of football impact reconstructions for head kinematics and finite element modeling. Clinical biomechanics 64, 82–89.

Subramaniam, D.R., Unnikrishnan, G., Sundaramurthy, A., Rubio, J.E., Kote, V.B., Reifman, J., 2021. The importance of modeling the human cerebral vasculature in blunt trauma. Biomedical engineering online 20, 1–19.

Sullivan, S., Eucker, S.A., Gabrieli, D., Bradfield, C., Coats, B., Maltese, M.R., Lee, J., Smith, C., Margulies, S.S., 2015. White matter tract-oriented deformation predicts traumatic axonal brain injury and reveals rotational direction-specific vulnerabilities. Biomechanics and modeling in mechanobiology 14, 877–896.

Tournier, J.-D., Calamante, F., Gadian, D.G., Connelly, A., 2004. Direct estimation of the fiber orientation density function from diffusion-weighted MRI data using spherical deconvolution. Neuroimage 23, 1176–1185.

Wang, T., Kleiven, S., Li, X., 2020. Electroosmosis based novel treatment approach for cerebral edema. IEEE Transactions on Biomedical Engineering 68, 2645–2653.

Wolf, J.A., Johnson, B.N., Johnson, V.E., Putt, M.E., Browne, K.D., Mietus, C.J., Brown, D.P., Wofford, K.L., Smith, D.H., Grady, M.S., 2017. Concussion induces hippocampal circuitry disruption in swine. Journal of neurotrauma 34, 2303–2314.

Wu, T., Alshareef, A., Giudice, J.S., Panzer, M.B., 2019. Explicit modeling of white matter axonal fiber tracts in a finite element brain model. Annals of biomedical engineering 47, 1908–1922.

Wu, T., Hajiaghamemar, M., Giudice, J.S., Alshareef, A., Margulies, S.S., Panzer, M.B., 2021a. Evaluation of tissue-level brain injury metrics using species-specific simulations. Journal of neurotrauma.

Wu, Y.-H., Rosset, S., Lee, T.-r., Dragunow, M., Park, T., Shim, V., 2021b. In Vitro Models of Traumatic Brain Injury: A Systematic Review. Journal of Neurotrauma.

Yang, K., Mao, H., 2019. Modelling of the Brain for Injury Simulation and Prevention, Biomechanics of the Brain. Springer.

Zhan, X., Li, Y., Liu, Y., Domel, A.G., Alizadeh, H.V., Zhou, Z., Cecchi, N.J., Raymond, S.J., Tiernan, S., Ruan, J., Barbat, S., Gevaert, O., Zeineh, M.M., Grant, G.A., Camarillo, D.B., 2021. Predictive Factors of Kinematics in Traumatic Brain Injury from Head Impacts Based on Statistical Interpretation. Annals of Biomedical Engineering.

Zhao, W., Cai, Y., Li, Z., Ji, S., 2017. Injury prediction and vulnerability assessment using strain and susceptibility measures of the deep white matter. Biomechanics and modeling in mechanobiology 16, 1709–1727.

Zhao, W., Ford, J.C., Flashman, L.A., McAllister, T.W., Ji, S., 2016. White matter injury susceptibility via fiber strain evaluation using whole-brain tractography. Journal of neurotrauma 33, 1834–1847.

Zhao, W., Ji, S., 2019a. Mesh convergence behavior and the effect of element integration of a human head injury model. Annals of biomedical engineering 47, 475–486.

Zhao, W., Ji, S., 2019b. White matter anisotropy for impact simulation and response sampling in traumatic brain injury. Journal of neurotrauma 36, 250–263.

Zhao, W., Ji, S., 2020. Incorporation of vasculature in a head injury model lowers local mechanical strains in dynamic impact. Journal of biomechanics 104, 109732.

Zhou, Z., Domel, A.G., Li, X., Grant, G., Kleiven, S., Camarillo, D., Zeineh, M., 2021a. White matter tract-oriented deformation is dependent on real-time axonal fiber orientation. Journal of neurotrauma 38, 1730–1745.

Zhou, Z., Jiang, B., Cao, L., Zhu, F., Mao, H., Yang, K.H., 2016. Numerical simulations of the 10-year-old head response in drop impacts and compression tests. computer methods and programs in biomedicine 131, 13–25.

Zhou, Z., Li, X., Domel, A.G., Dennis, E.L., Georgiadis, M., Liu, Y., Raymond, S.J., Grant, G., Kleiven, S., Camarillo, D., Zeineh, M., 2022. The Presence of the Temporal Horn Exacerbates the Vulnerability of Hippocampus During Head Impacts. Frontiers in Bioengineering and Biotechnology 10.

Zhou, Z., Li, X., Kleiven, S., 2019a. Biomechanics of acute subdural hematoma in the elderly: A fluid-structure interaction study. Journal of neurotrauma 36, 2099–2108.

Zhou, Z., Li, X., Kleiven, S., 2019b. Biomechanics of Periventricular Injury. Journal of Neurotrauma 37, 1074–1090.

Zhou, Z., Li, X., Kleiven, S., 2019c. Fluid–structure interaction simulation of the brain–skull interface for acute subdural haematoma prediction. Biomechanics and modeling in mechanobiology 18, 155–173.

Zhou, Z., Li, X., Kleiven, S., Hardy, W.N., 2019d. Brain strain from motion of sparse markers. Stapp car crash journal 63, 1–27.

Zhou, Z., Li, X., Kleiven, S., Shah, C.S., Hardy, W.N., 2018. A Reanalysis of Experimental Brain Strain Data: Implication for Finite Element Head Model Validation. Stapp car crash journal 62, 293–318.

Zhou, Z., Li, X., Liu, Y., Fahlstedt, M., Georgiadis, M., Zhan, X., Raymond, S.J., Grant, G., Kleiven, S., Camarillo, D., 2021b. Towards a comprehensive delineation of white matter tract-related deformation. Journal of neurotrauma 38, 3260–3278.

